# Identification of Genetic Suppressors for a Berardinelli-Seip Congenital Generalized Lipodystrophy Type 2 (BSCL2) Pathogenic Variant in *C. elegans*

**DOI:** 10.1101/2023.09.22.559059

**Authors:** Xiaofei Bai, Harold E. Smith, Andy Golden

**Affiliations:** Department of Biology, University of Florida, Gainesville, FL; Genetics Institute, University of Florida, Gainesville, FL; National Institute of Diabetes and Digestive and Kidney Diseases, National Institutes of Health, Bethesda, MD

## Abstract

Maintaining the metabolic homeostasis of fatty acids is crucial for human health. Excess fatty acids are stored in lipid droplets (LDs), the primary energy reservoir that helps regulate fat and lipid homeostasis in nearly all cell types. Seipin (BSCL2), a conserved endoplasmic reticulum protein, plays a critical role in LD biogenesis and regulating LD morphology. Pathogenic variants of seipin are associated with multiple human genetic diseases, including Berardinelli-Seip Congenital Generalized Lipodystrophy Type 2 (BSCL2). However, the cellular and molecular mechanisms by which dysfunctional seipin leads to these diseases remain unclear. To model BSCL2 disease, we generated an orthologous *BSCL2* pathogenic variant *seip-1(A185P)* using CRISPR/Cas9 genome editing in *Caenorhabditis elegans*. This variant led to severe developmental and cellular defects, including embryonic lethality, impaired eggshell formation, and abnormally enlarged LDs. We set out to identify genetic determinants that could suppress these defective phenotypes in the *seip-1(A185P)* mutant background. To this end, we conducted an unbiased chemical mutagenesis screen to identify genetic suppressors that restore embryonic viability in the *seip-1(A185P)* mutant background. A total of five suppressor lines were isolated and recovered from the screen. The defective phenotypes of *seip-1(A185P)*, including embryonic lethality and impaired eggshell formation, were significantly suppressed in each suppressor line. Two of the five suppressor lines also alleviated the enlarged LDs in the oocytes. We then mapped a suppressor candidate gene, *R05D3.2* (renamed as *lmbr-1*), which is an ortholog of human *LMBR1* (limb development membrane protein 1). The CRISPR/Cas9 edited *lmbr-1* suppressor alleles, *lmbr-1(Ser647Phe)* and *lmbr-1(Pro314Leu)*, both significantly suppressed embryonic lethality and defective eggshell formation in the *seip-1(A185P)* background. The newly identified suppressor lines offer valuable insights into potential genetic interactors and pathways that may regulate seipin in the lipodystrophy model.

## Introduction

Lipid droplets (LDs) are cellular energy reservoirs to maintain fat homeostasis and balance, ensuring proper lipid metabolism and healthy physiology. Failure to maintain the size and morphology of LD can lead to many diseases and disorders, including obesity, cancer, lipodystrophy, and neurodegenerative disorders [1–3]. Seipin is one of the primary LD scaffold proteins to govern LD size and may also regulate lipid biosynthesis and sorting to promote LD synthesis at the endoplasmic reticulum (ER) [4–6]. Clinical studies have identified multiple seipin pathogenic variants in patients with Berardinelli-Seip Congenital Lipodystrophy (BSCL), Silver syndrome, and Teratozoospermia syndrome [7–10]. Therefore, understanding the molecular and cellular mechanisms governing LD formation and the nature of LD scaffold proteins, such as seipin, are critical to developing therapeutic strategies for LD-associated diseases.

Proper lipid metabolism and transfer are critical to oogenesis, fertilization, embryogenesis, and other physiological processes, such as lifespan and locomotion, in *C. elegans*. Two lipid reservoirs, yolk granules and LDs, were identified in *C. elegans* oocytes and embryos to regulate lipid transfer and metabolism. However, the molecular and cellular mechanisms of how these two lipid structures contribute to healthy physiology remain unexplored. Our recent study established a *BSCL2* (Seipin) disease model for understanding seipin-associated disorders *in vivo* [11]. Three *seip-1* null alleles and an orthologous BSCL pathogenic variant allele *seip-1(A185P)* caused severe embryonic lethality and abnormally enlarged LDs in *C. elegans* oocytes and embryos. The embryonic lethality in the seipin mutants was correlatively linked to the impaired permeability barrier of the eggshell, a lipid-enriched extracellular matrix layer surrounding the embryo. Lipidomic studies showed that the C20 polyunsaturated fatty acid content was significantly reduced in the seipin mutants. The dietary supplementation of two types of linoleic acids restored embryonic lethality and extracellular eggshell formation in the seipin mutants. Intriguingly, the supplementation of the linoleic acids failed to rescue the abnormal/enlarged LD size and instead led to excessively enlarged LDs in oocytes and embryos [11], suggesting that proper eggshell and LD formation were independently regulated by seipin. However, the exact contributions of seipin during these two cellular events remain obscure.

The nematode model *Caenorhabditis elegans* has become an emerging system for understanding lipid and lipid metabolism diseases [12, 13]. At least ten lipodystrophy-associated genes have been identified, including phosphate acetyltransferase (*AGPAT2*), phosphoinositide-dependent serine-threonine protein kinase (*AKT2*), seipin (*BSCL2*), caveolin 1 (*CAV1*), and perilipin 1 (*PLIN1*) [14]. The conserved orthologs of all these lipodystrophy-associated genes are found in the *C. elegans* genome. Disturbing expression of the *C. elegans* orthologous genes led to various phenotypes, including embryonic lethality, sterility, lifespan alterations, and locomotion defects [11, 15–17]. These easily scorable phenotypes allowed geneticists to conduct forward genetic screens to identify novel genetic modifiers for LD biogenesis and lipid signaling pathways [18–20]. The development of rapid mapping strategies utilizing molecular inversion probes (MIP-MAP; [21]) combined with high-throughput whole-genome sequencing provides a fast and cost-effective method to pinpoint candidate mutations in a mutagenized genetic model.

To explore more specific roles that seipin plays during embryonic and LD biosynthesis, we focused on identifying its novel genetic determinants and pathways. Taking advantage of the embryonic lethality in the *seip-1(A185P)* mutant as a readout, we conducted an unbiased forward genetic screen to identify genetic modifiers that restore embryonic viability in the *seip-1(A185P)* mutant. Using MIP-MAP genomic mapping and the whole-genome sequencing technique, we identified two independent missense alleles of a target suppressor gene *R05D3.2*, which we renamed *lmbr-1*. We then generated those two putative suppressor alleles of *lmbr-1* gene, *lmbr-1(S647F)* and *lmbr-1(P314L)*, in the wild-type background. The *lmbr-1* missense mutants were then crossed back into the *seip-1(A185P)* mutant to assess their suppression. The homozygous *lmbr-1(S647F)* and *lmbr-1(P314L)* alleles significantly suppressed embryonic lethality and eggshell formation defects caused by *seip-1(A185P)*. *lmbr-1(S647F)* did not alleviate the enlarged LD size caused by *seip-1(A185P)*, while the other allele *lmbr-1(P314L)* partially suppressed enlarged LD size in the *seip-1(A185P)* background. In summary, we describe the results of the mutagenesis screen designed to identify regulators that genetically interact with a seipin BSCL2 pathogenic variant. The discovery of new suppressor candidates will shed light on the molecular mechanisms contributing to seipin-associated lipodystrophy and related physiological disorders.

## Results

### Berardinelli-Seip Congenital Generalized Lipodystrophy type 2 (BSCL2) pathogenic variant impaired eggshell formation and caused enlarged LD size but did not alter the sub-cellular localization of SEIP-1::mScarlet

Previously, we established a BSCL2 disease model by generating an orthologous pathogenic variant *seip-1(A185P)* in *C. elegans* [11], which corresponds to the orthologous Alanine212 in the human *BSCL2* gene. Using this disease model, we could functionally characterize the *seip-1(A185P)* allele *in vivo*. *seip-1(A185P)* caused penetrant embryonic lethality, defective eggshell formation, and abnormally enlarged LDs [11]. To determine whether this missense mutation disrupts the temporal and spatial expression pattern of SEIP-1, we generated the *seip-1(A185P)* allele by CRISPR/Cas9 genome editing into an endogenously tagged red fluorescent reporter strain, *seip-1::mScarlet* (Fig. 1A), which we designated *seip-1(A185P)::mScarlet*.

**Figure 1:**
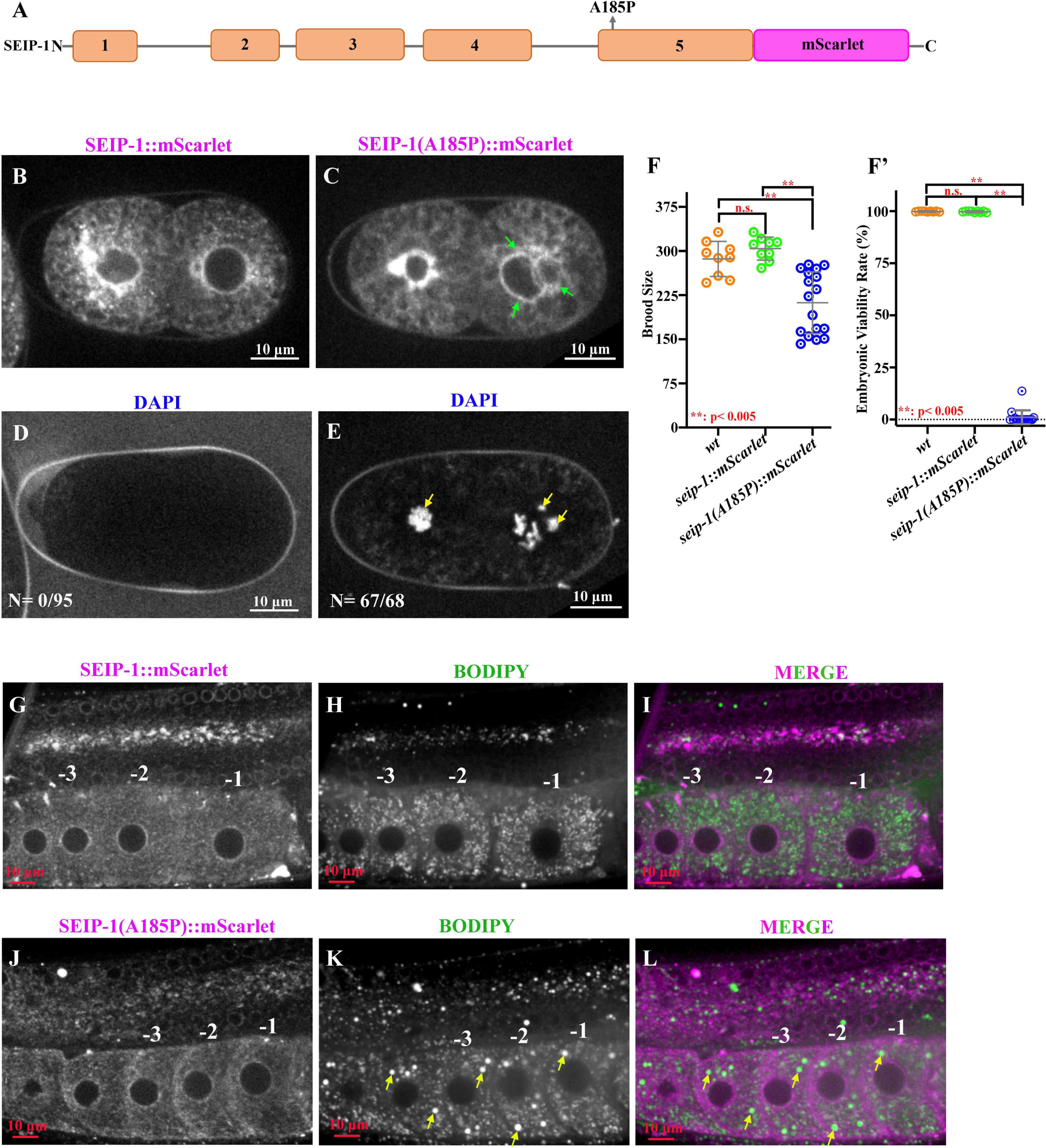
Functional analysis of the A185P mutation in a *seip-1∷mScarlet* reporter gene. (A) Schematic diagram of *seip-1(A185P)∷mScarlet*. (B-C) Visualization of mScarlet fluorescence to assess SEIP-1 localization. Green arrows indicate the presence of multiple nuclei in the posterior cell. (D-E) DAPI staining of embryos to assess eggshell permeability. Yellow arrows indicate the presence of stained chromatin. (F-F’) Brood size and embryonic viability of the mScarlet reporter genes. (G-L) SEIP-1 localization and lipid droplets in the adult germline. Representative images of animals containing the *seip-1∷mScarlet* (G-I) or *seip-1(A185P)∷mScarlet* (J-L) reporter as visualized by mScarlet protein fluorescence (G, J), the lipid dye BODIPY (H, K), or merged images (I, L). Yellow arrows indicate the presence of enlarged lipid droplets. Oocytes are numbered conventionally (−1, −2, −3) relative to the spermatheca, the site of fertilization. Scale bars are indicated in each panel.

Embryonic localization of *seip-1(A185P)∷mScarlet* was similar to the wild-type reporter, with enrichment around the nuclear envelope (Fig. 1B-C). This enrichment corresponds to the endoplasmic reticulum (ER), as *seip-1::mScarlet* was shown previously to co-localize with the ER marker SP12 [11]. However, the cytosolic distribution was more diffuse and less punctate in the *seip-1(A185P)∷mScarlet* mutant compared to the wild-type, suggesting a defect in ER structure or the association of SEIPIN with the ER. The most apparent difference was the presence of multiple nuclei in the *seip-1(A185P)∷mScarlet* strain (Figure 1C, green arrows). This defect has not been reported previously and likely contributes to the embryonic lethality observed in the *seip-1(A185P)* strain. *seip-1::mScarlet* was previously shown to express in additional *C. elegans* tissues, including the pharynx and epidermis, where the LDs are enriched [11, 22]; similar localization patterns were found in the *seip-1(A185P)::mScarlet* mutant (Fig. S1). Imaging of the adult germline (see following) also revealed that *seip-1(A185P)::mScarlet* distribution was normal (Fig. 1G, I-J, L). Taken together, results from the fluorescent reporter genes suggest that the *seip-1(A185P)* mutation may compromise protein function instead of altering the trafficking and cellular localization of SEIP-1 *in vivo*.

We performed a variety of functional assays to test this hypothesis. We used DAPI, a small molecule that can penetrate into embryos with an impaired eggshell permeability barrier, to assess eggshell integrity in the *seip-1(A185P)::mScarlet* mutant. In the *seip-1::mScarlet* control strain, no DAPI was staining detected in the cytosol, demonstrating that knock-in of *mScarlet* at the endogenous loci of *seip-1* did not disrupt eggshell integrity (N=0/95) (Fig. 1D). In contrast, zygotic chromatin labeled by DAPI was frequently observed in the fertilized embryos of the *seip-1(A185P)::mScarlet* mutant (N=67/68), which is indicative of impaired eggshell formation (Fig. 1E); we also observed multiple chromatin bodies (Fig 1E, yellow arrows) associated with the multi-nucleate phenotype, which is a novel observation in the *seip-1* mutants. Next, we assessed embryonic viability, as prior work indicated that the wild-type *seip-1::mScarlet* reporter did not affect survival [11]; in contrast, we found that *seip-1(A185P)::mScarlet* exhibited reduced brood size and embryonic lethality comparable to the *seip-1(A185P)* allele (Fig. 1F-F’). Finally, we examined LD size by staining animals with a lipophilic fluorescent probe BODIPY 493/503 in both *seip-1::mScarlet* control (Fig. 1G-I) and the *seip-1(A185P)::mScarlet* mutant (Fig. 1J-L). Enlarged LDs were observed in the oocytes on the proximal side of the gonad, which is adjacent to the spermatheca of the *seip-1(A185P)::mScarlet* animals (Fig. 1J-L). The defective phenotypes in the *seip-1(A185P)::mScarlet* mutant are consistent with other *seip-1* mutants characterized in the previous study [11]. We conclude that the primary impact of the A185P mutation is on protein function rather than localization, although a modest effect on the latter cannot be ruled out.

### EMS-based forward genetic screen to identify suppressors of *seip-1(A185P)*

To identify novel genetic modifiers that regulate seipin function, we conducted an ethyl methane sulfonate (EMS)-based forward genetic screen with the *seip-1(A185P)* mutant. The *seip-1(A185P)* allele caused a significantly reduced brood size, which includes both unhatched/dead embryos and hatched larvae and penetrant embryonic lethality (Fig. 2A-D) [11]. During the course of screen optimization, we determined that these defects were enhanced at a higher temperature (25°C) and that the embryonic lethality in *seip-1(A185P)* worsened when the animals were maintained at 25°C for one to two generations. Therefore, we performed the mutagenesis screen under this optimized temperature to isolate and recover suppressor lines of *seip-1(A185P)* that restored embryonic viability and produced viable larvae (Fig. 2B). The screen yielded a total of five independent suppressor lines from ∼64,000 haploid *C. elegans* genomes (∼32,000 mutagenized F1s). To generate the homozygous suppressor lines, embryonic lethality in each line was assessed for at least two generations, testing more than 20 animals per generation. Only the candidate lines displaying consistently elevated viability rates in each tested animal were collected and identified as the homozygous suppressor lines. Sanger sequencing confirmed the *seip-1(A185P)* allele in each suppressor line. The embryonic viability rate in each suppressor line was significantly increased when compared with *seip-1(A185P)* allele alone (Fig. 2D). Yet, the brood size in each suppressor line was reduced (Fig. 2C). Overall, our findings suggest that homozygous suppressor variants might alleviate the embryonic lethality caused by the *seip-1(A185P)* mutant. However, these variants could potentially have additional impacts on reproduction, such as causing a reduction in brood size. It is also plausible that other essential mutations in the suppressor background could contribute to the observed sub-fertility.

**Figure 2.**
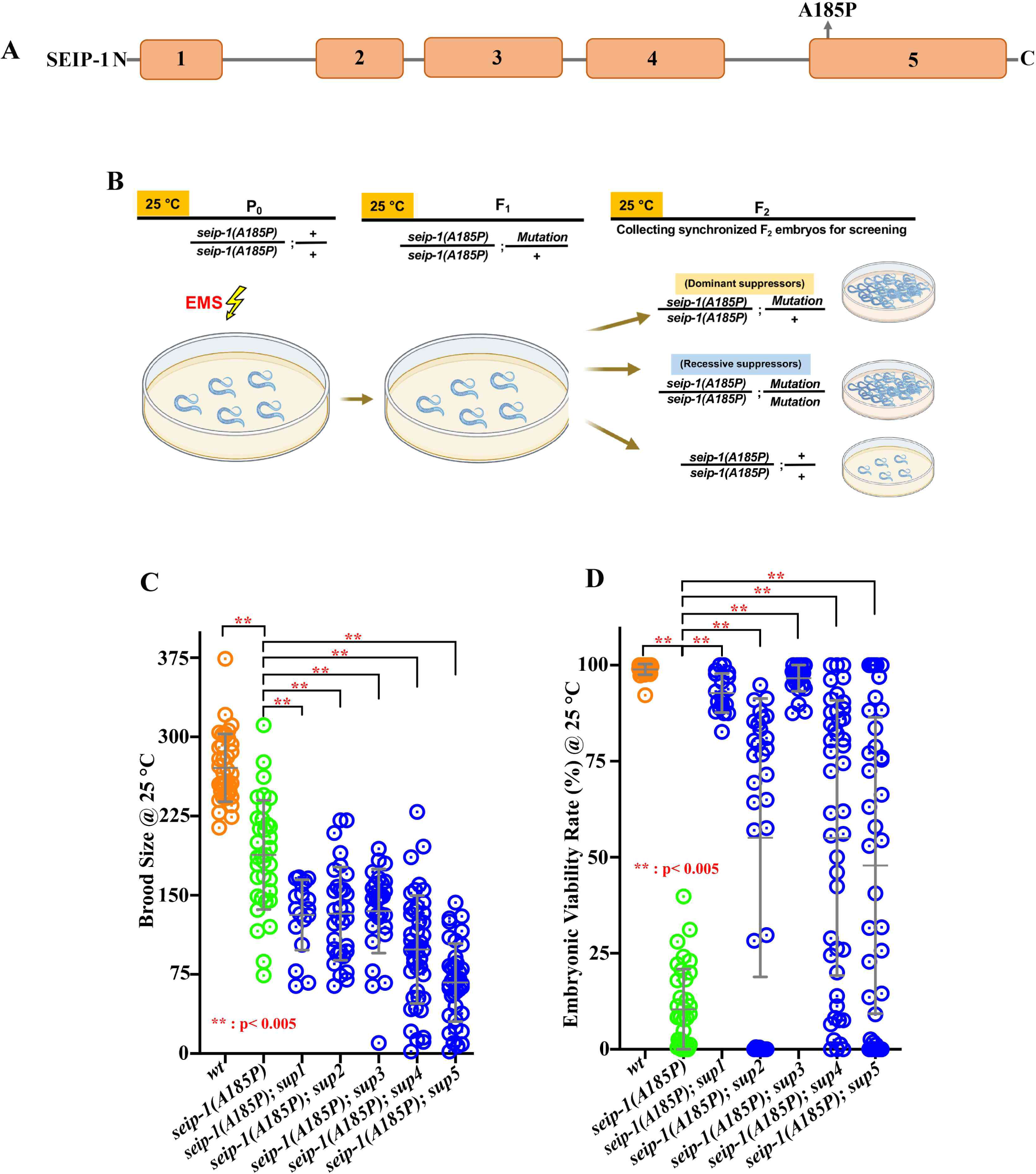
Forward genetic screen for suppressors of the embryonic lethality of a seipin patient allele. (A) Diagram of the location of the *seip-1(A185P)* mutation in the *seip-1* gene. (B) Strategy to identify genetic modifiers that restored embryonic viability in *seip-1(A185P)* mutant. (C) Total brood size of each isolated suppressor line, wild type, and *seip-1(A185P)* mutant over 60 hours post mid-L4. (D) The percentage of embryonic viability was significantly restored in each suppressor line when compared with *seip-1(A185P)* mutant only. Data are mean ± standard deviation. Statistical significance was determined using a One-Way ANOVA, ** p<0.005. The strategy graphic was generated with BioRender. com.

### The defective eggshell formation was significantly restored in each suppressor line

The previous study indicated that the embryonic lethality of the *seip-1(A185P)* mutant was correlatively linked with impaired eggshell [11]. To test whether the eggshell formation deficiency was restored in each suppressor line, we imaged the DAPI penetration in the early embryos. In the wild type, only the first polar body, located outside the permeability barrier (one layer of the eggshell) that blocks small molecule penetration, was stained by DAPI (Fig. 3A, C-E). In the *seip-1(A185P)* mutant, DAPI could penetrate the eggshell and stain the zygotic chromatin, indicating the defective permeability barrier (Fig. 3B, F-H). Following the DAPI imaging protocol, we tested over a hundred early embryos of each suppressor line. We found that DAPI penetration in each tested suppressor line was significantly reduced compared to *seip-1(A185P)* mutant (Fig. 3I). These observations suggested that the putative modifiers in the suppressor lines restore the proper formation of the permeability barrier. Not surprisingly, the suppressor lines *seip-1(A185P); sup1* and *seip-1(A185P); sup3,* which displayed the least DAPI penetration, correlated with the highest levels of embryonic viability (Fig. 2D and 3I), supporting our proposed hypothesis that the embryonic lethality in *seip-1(A185P)* mutant background is linked with the proper eggshell formation.

**Figure 3.**
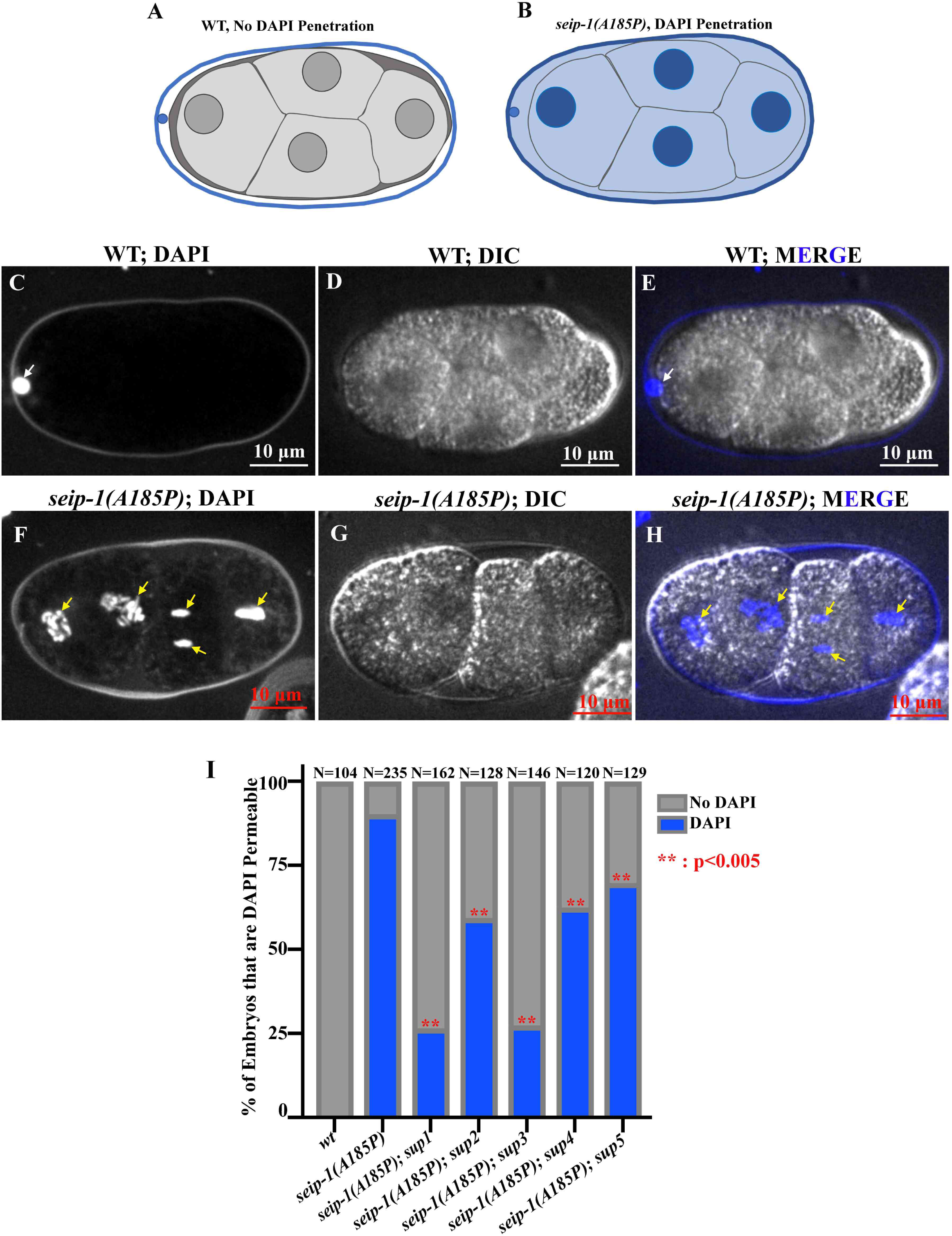
Defective eggshell formation was restored in each suppressor line. (A-B) Schematic diagram of DAPI staining assay in *C. elegans* embryos to assess eggshell permeability. (C-E) Representative images of the DAPI staining assay showed only the first polar body stained in wild-type (A, E, white arrow), whereas DAPI penetrated the eggshell to stain the zygotic chromatin in *seip-1(A185P)* mutation (F, H, yellow arrows). DIC images of embryos were shown in panels D and G. (I) Quantification of the embryos with DAPI staining in wild type, *seip-1(A185P)* only, and all suppressor lines. Statistical significance was determined using a Chi-square test. **: p< 0.005.

### Abnormal enlarged LD size was not correlatively rescued in each suppressor line of *seip-1(A185P)*

The canonical function of seipin is to maintain LD size and biosynthesis. We found that irregularly enlarged LDs were observed in the oocytes and embryos of the *seip-1(A185P)* mutant compared with wildtype [11](Fig. 4A-B, H). To test whether the abnormal LD size was alleviated in the suppressor lines, we imaged the oocytes where LDs were enriched by staining with a lipophilic BODIPY neutral fluorophore that binds to neutral lipids (Fig. 4A-G). The parameter of LD size was defined and quantified by measuring the number of LDs with diameters larger than 1.5 μm in the −1 to −3 oocytes as described in the previous study [11]. In wild type, the average LD size was less than 0.5 μm (Fig. 4A, H), but more than twenty enlarged LDs (> 1.5 μm) were readily stained in the −1 to −3 oocytes in the *seip-1(A185P)* mutant (Fig. 4B, H). The enlarged LD size (> 1.5 μm) was only alleviated in two of five suppressor lines, *seip-1(A185P); sup4* and *seip-1(A185P); sup5* (Fig. 4F-G, H). These two suppressor lines displayed the most rescued LD phenotypes. Still, they had the highest levels of DAPI penetration and embryonic lethality (Fig 2D and 3I), suggesting that genetic modifiers in each suppressor line may function somewhat independently in regulating eggshell formation, embryonic viability, and the maintenance of proper LD size in the oocyte. These observations are consistent with our previous observation that dietary supplementation of PUFAs significantly restored embryonic lethality and defective eggshell formation but enhanced the abnormal enlarged LD size [11]. This finding is also observed in a recent study indicating that the changes in phospholipid synthesis could suppress embryonic lethality but not affect LD size in a *seip-1* null mutant [34]. In summary, our data suggested that maintaining proper LD size primarily contributes to maintaining lipid balance and metabolism homeostasis but may only play a limited role in orchestrating eggshell formation.

**Figure 4.**
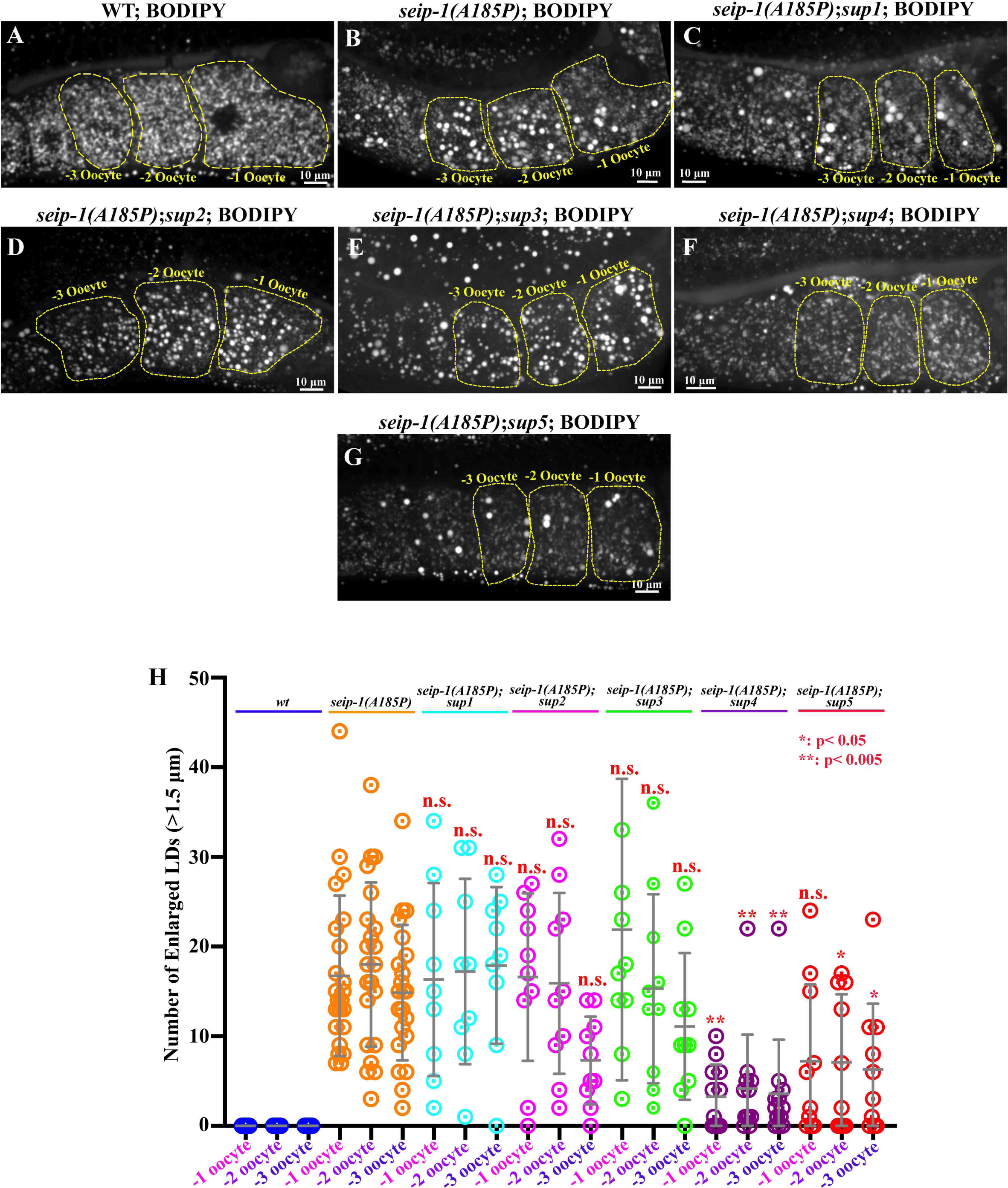
Enlarged LD was not correlatively restored in the suppressor lines. (A-G) BODIPY-stained LDs in the −1 to −3 oocytes of wild type (A), *seip-1(A185P)* only (B), and all five suppressor lines (C-G). (H) Quantification of the enlarged LDs (diameter >1.5 μm) inside the −1 to −3 oocytes of each stained genotype. Data are mean ± standard deviation. Statistical significance was determined using One-Way ANOVA. *: p< 0.05; **: p< 0.005. Scale bars are indicated in each panel.

### MIP-MAP mapping and suppressor allele validation

Using a fast and high-throughput genomic mapping strategy that involves molecular inversion probes (MIP-MAP) [21], we were able to identify the genetic modifiers in the suppressor line *seip-1; sup1* and *seip-1; sup4* (Fig. S2). We used a superficial wild-type strain, VC20019, with sufficient genomic diversity to provide ∼ 1 Mbp mapping resolution. To maintain the *seip-1(A185P)* allele during the mapping process, we generated *seip-1(A185P)* in the VC20019 background by CRISPR-mediated genome editing. After mating and propagation to select for animals that contained the suppressor, we constructed MIP-MAP and whole-genome libraries for next-generation sequencing. For two of the suppressor lines, *seip-1; sup1* and *seip-1; sup4*, we mapped the suppressors to overlapping genomic regions on chromosome III (*sup1,* 5.3-9.2 Mb, Fig. S2B; *sup-4,* 5.5-9.2 Mb, Fig. S2E). The mapping plots of the remaining suppressors either contained multiple gaps and peaks (*seip-1; sup3*) (Fig. S2D), indicative of multiple loci under selection, or no gaps (*seip-1; sup2* and *seip-1; sup5*) (Fig. S2C and F), consistent with dominant suppressors (Fig. S2). Therefore, we focused on suppressor lines *seip-1; sup1* and *seip-1; sup4.* After restricting the list of mutations to homozygous, protein-coding variants in the mapped regions and comparing the lists, we identified a single candidate modifier gene, *R05D3.2*, that contained different missense mutations in the two suppressor lines: *R05D3.2(S647F)* in *seip-1; sup1* and *R05D3.2(P314L)* in *seip-1; sup4 (*Fig. 5A). *R05D3.2* is an ortholog of human *LMBR1* (limb development membrane protein 1), and we renamed the *C. elegans R05D3.2* to *lmbr-1*.

**Figure 5.**
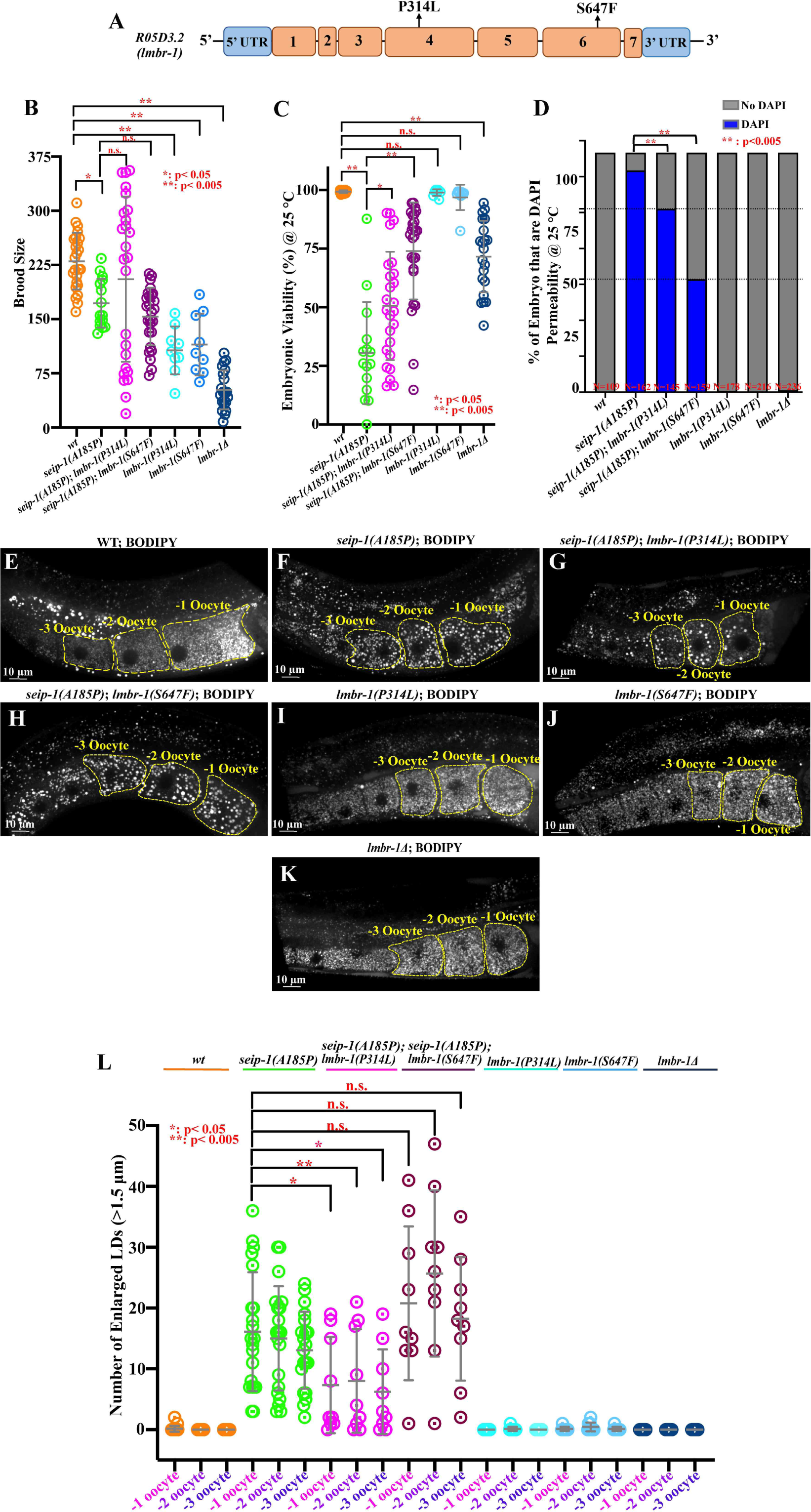
Identification of the genetic modifiers by MIP-MAP genome mapping and whole-genome sequencing. (A) Schematic diagram of two suppressor alleles of *lmbr-1* were generated in the wildtype background. (B-C) Total brood size and embryonic viability of each isolated wild type, *seip-1(A185P)*, *lmbr-1* suppressor alleles, *lmbr-1* null mutant, and double mutants of *seip-1(A185P)* and *lmbr-1* alleles. (D) DAPI penetration in the early embryos of each tested mutant and wild type. (E-K) Representative images of the BODIPY-stained LDs in the −1 to −3 oocytes of *lmbr-1(P314L), lmbr-1(S647F), lmbr-1Δ*, *lmbr-1(P314L); seip-1(A185P),* and *lmbr-1(S647F); seip-1(A185P)* double mutants. (L) Quantification of the enlarged LDs (diameter >1.5 μm) in the indicated genotype backgrounds. Statistical significance was determined using a Chi-square test, **: p< 0.005 in panel D, and One-Way ANOVA, *: p< 0.05, **: p<0.005 in panel L.

To validate *lmbr-1* as the suppressor candidate, we regenerated the two putative suppressor alleles of the *lmbr-1* gene, *lmbr-1(P314L)* and *lmbr-1(S647F)*, in the wild-type background by CRISPR/Cas9-mediated genome editing (Fig. 5A). Both *lmbr-1(P314L)* and *lmbr-1(S647F)* mutations alone caused reduced brood size compared to wild type (Fig. 5B), however, there were no embryonic lethality or eggshell formation defects observed in the *lmbr-1* missense mutants (Fig. 5C-D). To further characterize the role of *lmbr-1* in *C. elegans*, we also generated a null mutation, *lmbr-1Δ*, which bears a full-length deletion of the *lmbr-1* gene. Compared to the P314L and S647F alleles, the homozygous *lmbr-1Δ* mutant caused smaller brood size and ∼25% embryonic lethality (Fig. 5B-C); this result indicates that *lmbr-1* is critical to the *C. elegans* reproduction and embryogenesis, and suggests that the missense mutations are not loss-of-function alleles. Additionally, we did not observe any eggshell defects in the *lmbr-1Δ* mutant (Fig. 5D), suggesting that the embryonic lethality caused by *lmbr-1Δ* is not associated with the integrity of eggshell. To further investigate the expression pattern of *lmbr-1* in *C. elegans*, we successfully knocked-in a fluorescent reporter *gfp* at the N-terminus loci of the *lmbr-1* gene. However, we could not detect any visible fluorescent signal of GFP::LMBR-1, likely due to the low expression level of *lmbr-1* in *C. elegans*.

We then crossed the newly generated *lmbr-1(P314L)* and *lmbr-1(S647F)* alleles into the *seip-1(A185P)* background to test their suppression in the context of embryonic lethality, eggshell formation, and enlarged LD size. Both *lmbr-1(P314L)* and *lmbr-1(S647F)* significantly suppressed embryonic lethality and the defective eggshell formation but not affect the brood size compared to *seip-1(A185P)* alone (Fig. 5B-D). Additionally, all *lmbr-1* mutants, including *lmbr-1(P314L)*, *lmbr-1(S647F)*, and *lmbr-1Δ* alone did not affect LD size in the −1 to −3 oocytes compared to wild type (Fig. 5E, I-L). Intriguingly, the *lmbr-1(P314L)* allele partially suppressed the irregularly enlarged LD size in the −1 to −3 oocytes of the *seip-1(A185P)* mutant (Fig. 5F, G, and L). This finding is consistent with the alleviated enlarged LDs phenotype in the original suppressor line *seip-1(A185P); sup4*, which contains the *lmbr-1(P314L)* allele. The other allele *lmbr-1(S647F)*, like the *seip-1(A185P); sup1* strain from which it was identified, did not affect the enlarged LDs compared to the *seip-1(A185P)* mutant (Fig. 5F, H, and L). Overall, the CRISPR-edited *lmbr-1* alleles displayed identical suppression of embryonic lethality, defective eggshell formation, and enlarged LD phenotypes that were observed in the original suppressor lines. Additionally, different *lmbr-1* suppressor alleles, such as P314L and S647F, may contribute independently to regulating embryonic lethality/defective eggshell integrity and enlarged LDs size.

## Discussion

The primary objective of the forward genetic screen in the orthologous lipodystrophy *seip-1(A185P)* mutant background was to uncover novel genetic determinants and regulators of seipin. We sought to identify and characterize the genetic modifiers capable of restoring cellular and developmental deficiencies in the *seip-1(A185P)* mutant background. In the course of this study, we successfully isolated five distinct suppressor lines. Furthermore, we refined the MIP-MAP strategy by introducing the pathogenic variant into the mapping strain using CRISPR/Cas9 editing. This strategic innovation significantly streamlined the mapping process and reduced both labor and time required for identifying potential modifier candidates. During the trial screen, we identified two suppressor alleles, including *lmbr-1(P314L)* and *lmbr-1(S647F)*, within the newly implicated genetic determinant, *lmbr-1*. These alleles demonstrated a remarkable ability to suppress embryonic lethality and defective eggshell formation resulting from the *seip-1(A185P)* mutation.

### LMBR-1 and its suppression in *seip-1(A185P)* mutant

The *C. elegans* gene *R05D3.2* is the only ortholog of human *LMBR1* (limb development membrane protein 1) and *LMBR1L* (LMBR1-like membrane protein). Both LMBR1 and LMBR1L exhibit high conservation, sharing 60% identity in their amino acid sequence. A distinguishing feature found among all LMBR1 proteins is the presence of multiple transmembrane segments. These LMBR1 proteins are predicted to express at the ER membrane, similar to seipin expression [26]. Notably, the LMBR1L protein is also known as lipocalin-1 interacting membrane receptor (LIMR) [27], which interacts with lipocalin-1, an extracellular protein responsible for binding hydrophobic ligands, fatty acids, and phospholipid, thereby facilitating their internalization and subsequent degradation [28–30].

Therefore, the *lmbr-1* suppressor mutations are plausible candidates to influence lipid transfer or lipid metabolism, potentially acting to restore the disturbed lipid balance within the *seip-1(A185P)* mutant. Consequently, these mutations may contribute to the suppression of embryonic lethality and other defective phenotypes. Further investigations will be directed toward a better understanding of the underlying molecular mechanism driving this suppression. Particularly, we aim to characterize the intricate interplay between lipid metabolism and the functional coordination between seipin and *lmbr-1*.

In the scope of this study, we also characterized the functional contribution of *lmbr-1* by generating multiple mutants, including two missense suppressor alleles and a *lmbr-1* null mutant bearing a full-length deletion. All tested *lmbr-1* mutants displayed a reduction in brood size, and the null mutant exhibited about 30% of embryonic lethality. Interestingly, prior research has shed light on the role of LMBR1L as a pivotal player in cell signaling regulation. Specifically, it is known to modulate Wnt/Catenin and BMP signaling pathways in contexts of lymphocyte development and *Drosophila* oogenesis [26, 31]. Collectively, these findings underscore that *lmbr-1* may serve diverse functional roles which significantly contribute to the intricate dynamics of *C. elegans* reproduction and embryonic development. The observations also serve as an explanation for the reduced brood size in the original *seip-1* suppressor lines.

Lastly, the CRISPR-generated *lmbr-1* mutations showed a relatively low level of suppression of DAPI penetration compared to the EMS-derived strains, suggesting that additional modifiers likely await discovery. Meanwhile, we could not exclude the possibility that other cellular signaling pathways may be involved in regulating lipid droplet, embryogenesis, and eggshell formation in the seipin mutants. *lmbr-1* suppressor alleles may also be involved in other signaling pathways to coordinate the suppression of the defects.

### Congenital lipodystrophy disease modeling and future direction

In this study, we observed the emergence of a multi-nucleate and defective nuclear envelope formation, which appears to play a pivotal role in the observed lethality of the *seip-1(A185P)* mutant. Furthermore, similar nuclear envelope phenotypes were also associated with lipodystrophies, such as partial lipodystrophy of the Dunnigan type (FPLD2) [46]. Thus, cellular-level similarities could be relevant to human diseases, even though *C. elegans* lacks adipose tissues, which show significant phenotypes in human lipodystrophy conditions.

Our study also introduces an accessible *in vivo* system for investigating seipin in the *C. elegans* germline tissue, which is enriched in LDs and necessitates the transfer of lipids and fatty acids between somatic tissue during ovulation and fertilization. The mechanisms involving lipid transportation and metabolic regulation of these structures are likely conserved between humans and *C. elegans*. Moreover, the observed embryonic lethality in *seip-1(A185P)* mutants has allowed us to screen genetic antagonists specific to *seip-1* patient-specific alleles *in vivo*. For future studies, we intend to identify the novel genetic modifiers in the other three suppressor lines using the alternative mapping crosses more suitable for polygenic or dominant mutations. Our suitable paradigm of embryonic lethality in seipin mutants should allow us to continue forward screens to identify molecular pathways or determinants to suppress or reverse the defects. The findings in our genetic study may provide insights for future therapeutic targets. A successful reproductive cycle is precisely coordinated by lipid or modified lipid signaling to fertilization, oocyte growth, meiotic progression, and gonadal muscle contraction in *C. elegans*. The overarching theme of our study is to delineate the signaling mechanism of seipin in coordinating these physiological processes. Understanding these signaling pathways involving seipin should advance our understanding of lipid biology in other organisms, including humans.

## Materials and Methods

### Strain Maintenance

Strains used in this study were maintained on MYOB plates as previously described at 20°C or 25°C when screening the suppressors [32]. Detailed information of the strains are listed below: N2 Bristol (wild-type); AG429 *seip-1(av169[A185P]) V*. CRISPR/Cas9 Edit; AG444 *seip-1(av169[seip-1::mScarlet]) V*. CRISPR/Cas9 Edit; AG666 *seip-1(A185P) V; MIP-MAP*. CRISPR/Cas9 Edit; AG685 *seip-1(av304[seip-1(A185P)::mScarlet]) V*. CRISPR/Cas9 Edit; AG670 *seip-1(av169[A185P]) V; sup1;* AG671 *seip-1(av169[A185P]) V; sup2;* AG686 *seip-1(av169[A185P]) V; sup3;* AG687 *seip-1(av169[A185P]) V; sup4;* AG688 *seip-1(av169[A185P]) V; sup5;* AG743 *lmbr-1(av288[P314L]) III*; AG746 *lmbr-1(av291[S647F]) III;* AG750 *lmbr-1(av293[lmbr-1Δ] III*; AG751 *seip-1(av294) V*; AG755 *seip-1(av294) V*; *lmbr-1(av288[P314L]) III*; AG756 *seip-1(av294) V*; *lmbr-1(av291[S647F]) III*.

### EMS Suppressor Screen

*seip-1(A185P)* early L4 hermaphrodites were washed three times in M9 buffer and soaked in 48 mM Ethyl methane sulfonate (EMS) solution for four hours at room temperature. The EMS-treated animals were washed three times in M9 buffer and were then transferred to a fresh 100 mm MYOB plate with OP50 on one side. The animals were allowed to recover up to 4 hours before being picked to 100 mm MYOB plates with fresh OP50. Only the recovered animals that were able to crawl across the plates to the OP50 food were transferred to the fresh plates. A total of 15 MYOB plates with 10-15 mid-L4 (P0s) on each were incubated at 20°C. Gravid F1 adult progeny (∼32,000) were collected for synchronizing the F2 population using hypochlorite treatment. The F2 embryos were shaken in a glass flask with M9 buffer overnight at 20°C, the hatched larvae were shifted to 25°C, and their F3 progeny were screened for viable larvae. A total of five suppressor lines were isolated from the screen. The males of *seip-1(A185P^MM^)* were mated with the homozygous hermaphrodites of each suppressor line (Fig. S2A). We carefully pooled F2 progeny from the cross and allowed the F2 population to expand for over ten generations for MIP-MAP analysis, which is sufficient to distribute the MIP-MAP molecular probes into the suppressor line background and provides a high molecular resolution to identify the mutation regions (Fig. S2A). Theoretically, we should be able to narrow down the target regions bearing the putative modifiers by reading the frequency of MIP-MAP probes. Since the flanking regions of the modifiers originated from a wildtype background, the occurrence frequency of the MIP-MAP probes at the nearby loci would drop to nearly zero [21].

### Microscope and Imaging Analysis

For imaging SEIP-1::mScarlet and SEIP-1(A185P)::mScarlet expression, animals were immobilized on 7% agar pads with anesthetic (0.1% tricaine and 0.01% tetramisole in M9 buffer). DIC and mScarlet image acquisition were conducted using a Nikon 60×1.2 NA water objective with 1 μm *z*-step size; 20-25 *z* planes were captured. The image was performed on a scanning disk confocal system, including a Yokogawa CSU-X1 confocal scanner unit, a Nikon 60 x 1.2 NA water objective, and a Photometrics Prime 95B EMCCD camera. The images were analyzed by Nikon’s NIS imaging software and ImageJ/FIJI Bio-formats plugin (National Institutes of Health) [33, 34].

### DAPI staining of embryos

Gravid hermaphrodites were subject to three washes in M9 buffer and subsequently dissected using 23 G×3/4″ needles. Embryos at various developmental stages were then transferred into a handing drop chamber which was pre-filled with blastomere culture medium (BCM) as described in the previous study [23]. The BCM was prepared freshly, containing 10 ug/ml DAP for staining. Prior to imaging, the hanging drop chamber was sealed with molten Vaseline. Image acquisition was performed using a Nikon 60 x 1.2 NA water objective with a z-step size of 1 μm.

### BODIPY Staining

The lipophilic molecule BODIPY 493/503 (Invitrogen, D3922) was dissolved in 100% DMSO to 1 mg/ml. The working solution of BODIPY was diluted by M9 buffer to 6.7 μg/ml BODIPY (the final concentration of DMSO was 0.8%). The tested animals were picked and incubated in 100 μl of 6.7 μg/ml BODIPY for 30 minutes and soaked in M9 buffer for 5-10 minutes until the animals were hydrated and started moving. The recovered animals were then anesthetized with 0.1% tricaine and 0.01% tetramisole in M9 buffer for 15 to 30 minutes before being transferred to 7% agarose pads for imaging. The diameter of LD was quantified in each oocyte by FIJI ImageJ.

### CRISPR Design, Experiments, and Sequence information

The sequence information of CRISPR design is listed in Table 1. We followed the optimized CRISPR/Cas9 editing protocol that was used in our previous studies [11]. The gene-specific guide RNAs were designed with the help of a guide RNA design checker from Integrated DNA Technologies (www.idtdna.com) and were ordered as 4 nmol products from Horizon Discovery (www.dharmacon.horizondiscovery.com), along with tracRNA. Repair template design followed the standard protocols [35]. Young gravid animals (∼20) were injected with the prepared CRISPR/Cas9 injection mix as described in the literature [35, 36]. The Cas9 protein was purchased from PNA Bio (CP01). All homozygous animals edited by CRISPR/Cas9 were validated by Sanger sequencing. The detailed sequence information for the repair template and guide RNAs are listed in Table 2.

**Table 1.**
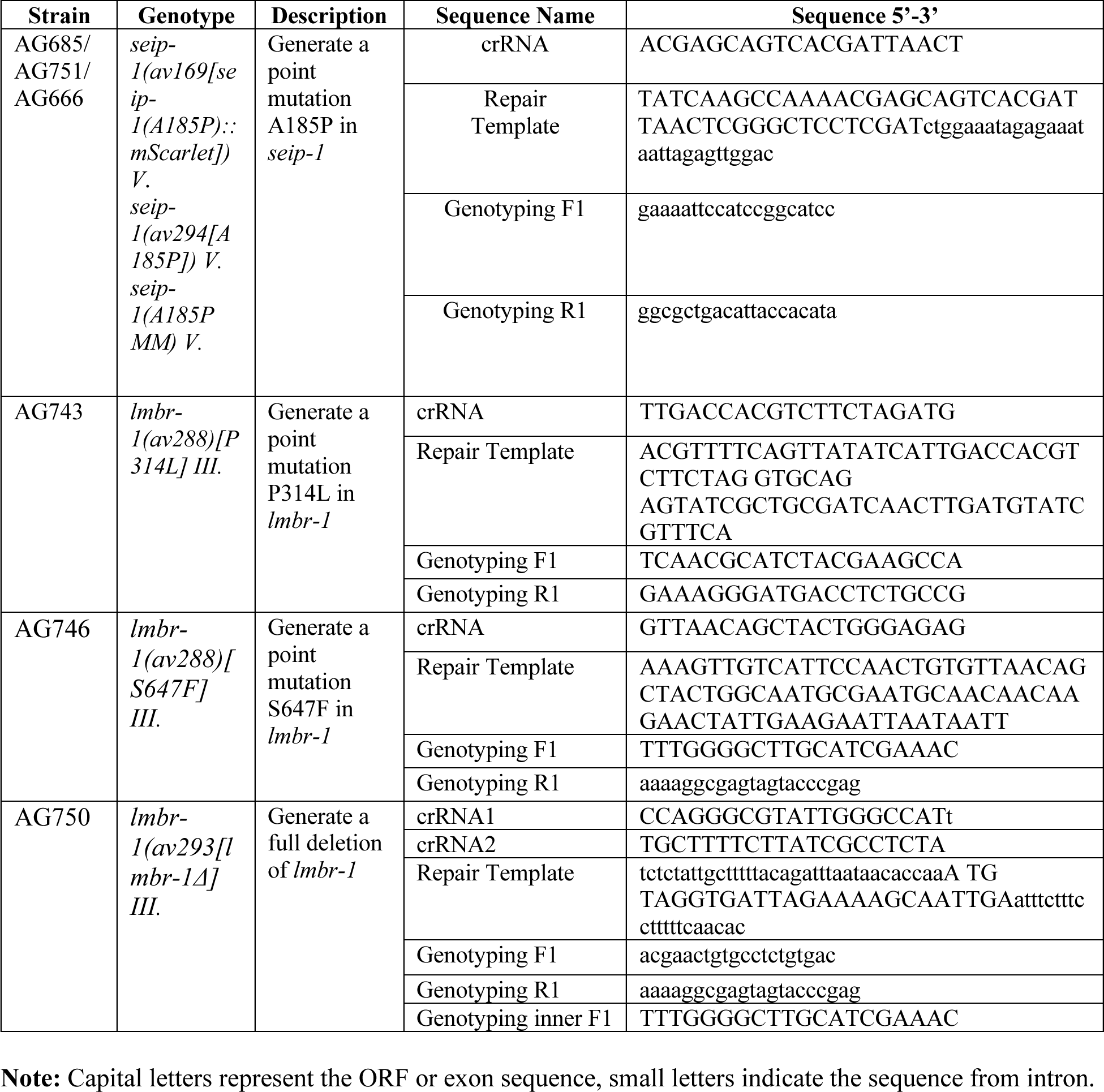
List of the sequence for the CRISPR design.

### MIP-MAP and Data analysis

Candidate mutations (defined as novel, homozygous, and nonsynonymous) were identified by whole-genome sequencing (WGS) as described previously [37]. Briefly, sequencing libraries were constructed by Invitrogen Pure Link Genomic DNA Mini Kit (Ref # K1820-01) with genomic DNA from homozygous suppressor-bearing strains. The WGS libraries were pooled and sequenced on a HiSeq 3000 instrument (Illumina, San Diego, CA) to at least 20-fold coverage. Variants were identified with a pipeline of BBMap [41], SAMtools [38], FreeBayes [42], and ANNOVAR [43]. Mapping loci for suppressors were identified using molecular inversion probes (MIPs) to single-nucleotide polymorphisms (SNPs) as described previously [44]. Briefly, suppressor-bearing strains were mated to SNP mapping strain VC20019 [39], engineered via CRISPR to contain the *seip-1(A185P)* mutation. F1 cross-progeny was allowed to self-fertilize, and a minimum of 50 homozygous F2 progeny were pooled for the construction of MIP libraries. SNP allele frequencies were determined using a custom script and plotted with R [45] to delimit the mapping interval.

## Acknowledgments

We thank past and present members of the Golden Lab for insightful discussions during the screening and preparation of the manuscript. We thank all members of the NIH Worm Club and the Baltimore Worm Club, Drs. Chao-Wen Wang (SINICA, Taiwan) and Will Printz (UT Southwestern Medical Center) for providing critical comments on the manuscript and our seipin project. The N2 strain was provided by the CGC, which is funded by the NIH Office of Research Infrastructure Programs (P40 OD010440). This work is dedicated in memory of Dr. Andy Golden, whose invaluable support greatly contributed to the success of this project. He will be deeply missed.

## Author’s Contributions

X.F.B., H.S., and A.G. designed the screen and methods for mutant characterization; X.F.B. performed the screen; H.E.S. performed MIP-MAP and WGS to identify the *lmbr-1* alleles. X.F.B., H.E.S., and A.G. wrote and edited the manuscript.

---

**Supplemental Figure 1.**
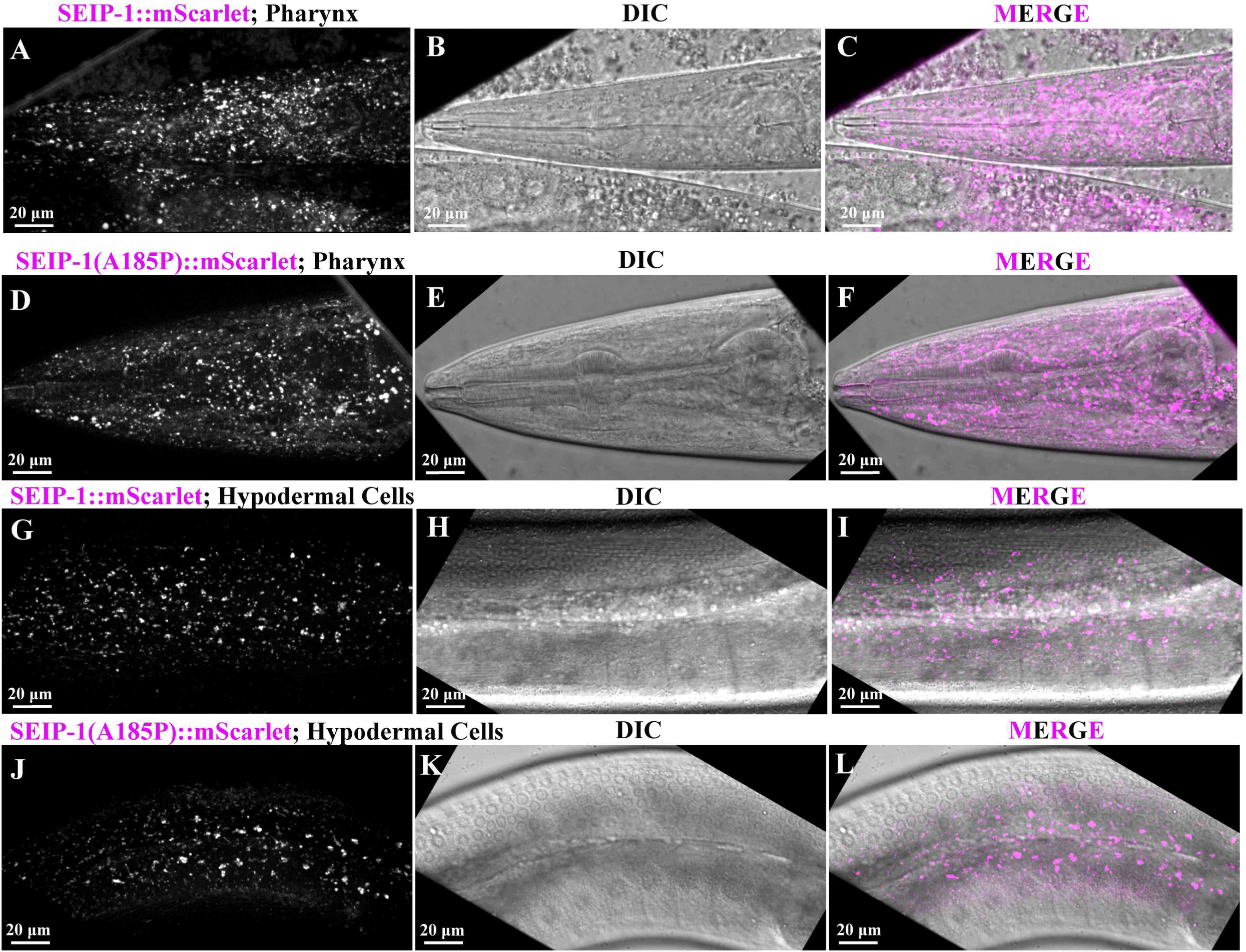
*seip-1(A185P)::mScarlet* does not disrupt the cellular localization of SEIP-1::mScarlet. (A-L) *seip-1(A185P)::mScarlet* (magenta, D, F, J, and L) presents an identical expression pattern as *seip-1::mScarlet* only (magenta, A, C, G, and I) in a variety of cell types, including pharynx (A-F) and hypodermal cells (G-L). DIC images are shown in panels B, E, H, and K. Scale bars are indicated in each panel.

**Supplemental Figure 2.**
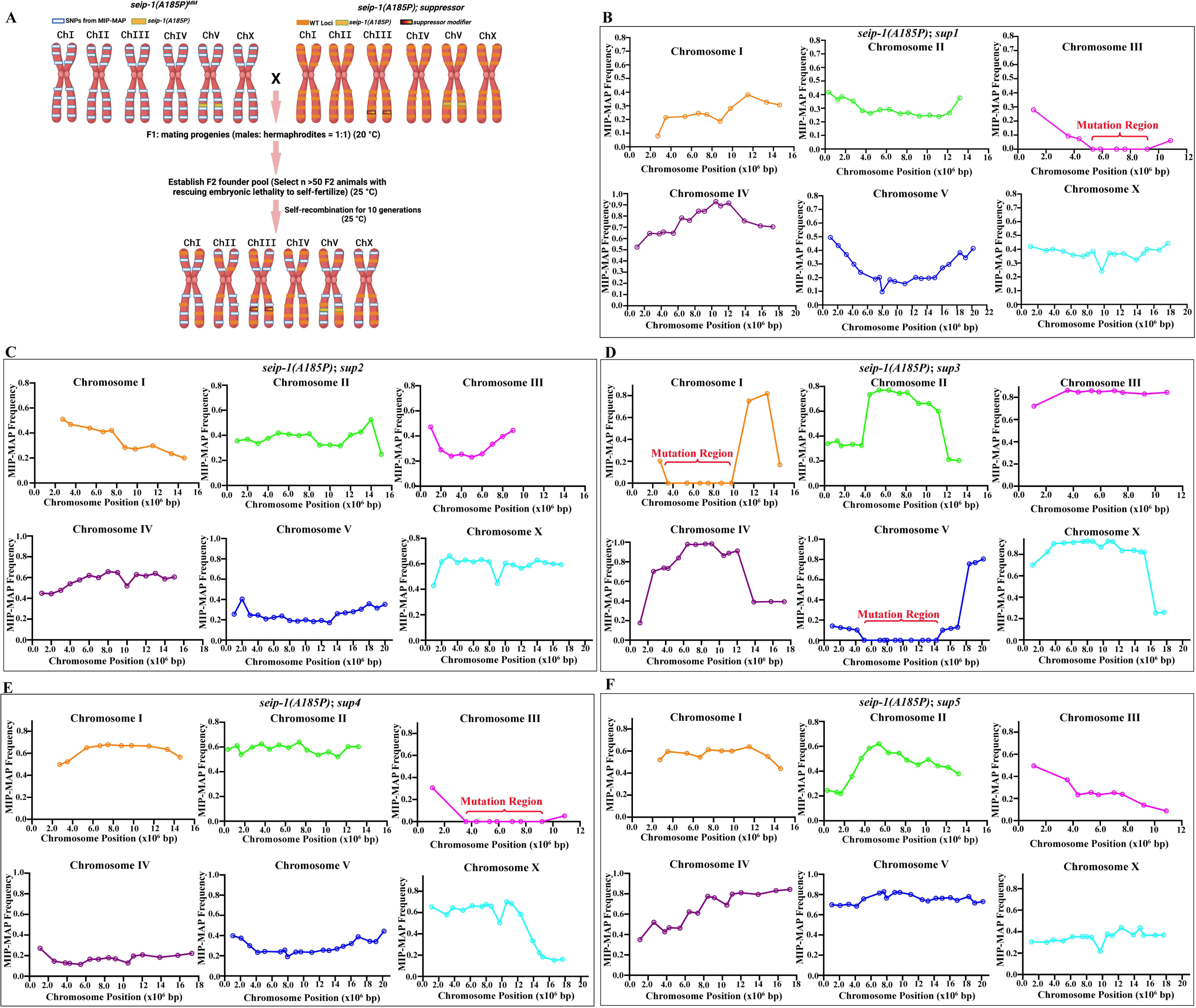
Mapping of *seip-1(A185P)* suppressors via VC20019 and MIP-MAP sequencing. (A) Diagram of the MIP-MAP workflow to map the SNVs in the suppressor line. (B-F) The read frequency of the VC20019-specific SNVs across the genome of the pooled F2 progeny from *seip-1(A185P); sup1-5* and *seip-1(A185PMM)* cross. (B, E) An identical candidate mutation-associated interval was identified on Chromosome III in *seip-1(A185P); sup1*, and *seip-1(A185P); sup4*. The strategy graphic was generated with BioRender. com.

